# Development of molecular markers for determining continental origin of wood from white oaks (*Quercus* L. sect. *Quercus*)

**DOI:** 10.1101/038562

**Authors:** H. Schroeder, R. Cronn, Y. Yanbaev, T. Jennings, M. Mader, B. Degen, B. Kersten

## Abstract

To detect and avoid illegal logging of valuable tree species, identification methods for the origin of timber are necessary. We used next-generation sequencing to identify chloroplast genome regions that differentiate the origin of white oaks from the three continents; Asia, Europe, and North America. By using the chloroplast genome of Asian Q. mongolica as a reference, we identified 861 variant sites (672 single nucleotide polymorphisms (SNPs); 189 insertion/deletion (indel) polymorphism) from representative species of three continents (Q. mongolica from Asia; Q. petraea and Q. robur from Europe; Q. alba from North America), and we identified additional chloroplast polymorphisms in pools of 20 individuals each from Q. mongolica (789 variant sites) and Q. robur (346 variant sites). Genome sequences were screened for insertion/deletion (indel) polymorphisms to develop markers that identify continental origin of oak species, and that can be easily evaluated using a variety of detection methods. We identified five indel and one SNP that reliably identify continent-of-origin, based on evaluations of up to 1078 individuals representing 13 white oak species and three continents. Due to the size of length polymorphisms revealed, this marker set can be visualized using capillary electrophoresis or high resolution gel (acrylamide or agarose) electrophoresis. With these markers, we provide the wood trading market with an instrument to comply with the U.S. and European laws that require timber companies to avoid the trade of illegally harvested timber.

## Introduction

Illegal logging is a serious issue not only for tropical rainforests and tropical trees, but it is also a concern for tree species in temperate latitude forests. White oaks from the genus *Quercus* sect. *Quercus* (Fagaceae) provide a relevant example of illegal logging in a temperate zone tree, and they highlight the challenge facing importers and regulatory agencies responsible for validating the taxonomic and geographic sources of timber products. White oaks account for a significant percentage of the hardwood flooring and furniture trade in Europe and the USA, and they represent one of the most important hardwoods in terms of logs and lumber exports from these regions. The most important trade woods of white oaks derive from the European species *Quercus robur* L. and *Q. petraea* (Mattuschka) Liebl., the CITES Appendix III-protected *Q. mongolica* Fisch. Ex Ledeb. native to East-Asia, and North American oaks, such as *Q. alba* L. and *Q. macrocarpa* Michx. (Cassens 2007). Non-governmental organizations such as the Environmental Investigation Agency (http://eia-global.org/news-media/liquidating-the-forests) have documented increases in the rate of illegal logging for white oak wood, especially in the Russian Far East region. These activities increase the likelihood that international wood trading companies will market illegally harvested wood, an activity that is banned by the U.S. Lacey Act amendment of 2008 and the European Union timber regulation of 2010. Violation of these regulations can result in fines, forfeiture of wood, and additional payments, as was recently demonstrated with improperly documented shipments of white oak flooring in the United States (US Department of Justice, 2015). Under these laws, timber companies are responsible for avoiding the trade of illegally harvested timber, and they are obligated to declare the species name and geographic origin of traded timber in order to reduce the risk that traded timber originated from illegal logging (Dormontt *et al.* 2015).

The increased attention to illegal logging has led to an increased demand for methods that can be used to provide precise species identification and geographic origin verification. Wood anatomical methods are widely used for tree species identification (Dormontt *et al.* 2015), but these methods cannot discriminate white oak species, nor identify geographic origin of oaks generally. Over the last decade, worldwide programs have been established using the potential of DNA as universal tool for identifying organisms (Barcode of Life www.barcodeoflife.org, Hollingsworth *et al.* 2009). In plants, the success of barcoding is highly dependent on several factors, including magnitude of primary divergence, frequency of secondary contact, and mutation rate of the DNA region (Hollingsworth *et al.* 2011), so the choice of suitable barcode regions in plants can be difficult (Chase *et al.* 2005; Newmaster *et al.* 2006; Kress & Erickson 2008). Barcoding efforts in plants have focused on chloroplast genomes due their simple pattern of (typically) uniparental inheritance, low effective population size, and useful variation at the scale of geography and taxonomy across a wide range of species (e.g. Kress & Erickson 2007; Taberlet *et al.* 2007; Lahaye *et al.* 2008; Janzen *et al.* 2009; Huang *et al.* 2015). With advances in next-generation sequencing, chloroplast genomes are affordable to sequence in their entirety by ‘skimming’ methods (Straub *et al.* 2012), and whole genome analysis can reveal substantial variation, even in unexpected genomic regions that are included in traditional barcoding efforts (e.g., Parks *et al.* 2009, 2012). DNA barcoding already has proven to be appropriate for revealing illegal trading (e.g. Goncalves *et al.* 2015; Pappalardo & Ferrito 2015), and it is increasingly used to identify plant species in commercial trade (e.g., Handy *et al.* 2011).

The aim of this study is to use chloroplast-genome scale information to develop a cost-efficient, easy-to-use assay that allows the identification of the geographic origin of white oak wood products to hemisphere (Old World vs. New World) and continent (Asia; Europe; North America) to support regulatory and commercial efforts to detect illegal logging of *Q. mongolica.*

## Material and Methods

### Plant material

For next-generation sequencing, we sequenced a single oak individual from four species and three continents to produce chloroplast genome references; included are *Q. mongolica* from Asia (sample QUMO5_CH_1; China), *Q. petraea* from Europe (sample QUPE2_PO_1; Poland), *Q. robur* from Europe (sample QURO2_SVT6; Germany), and *Q. alba* from North America (sample QUAL_VT_1; USA). To develop a panel of polymorphisms for Asia and Europe, we used next-generation sequencing to screen two pooled DNA samples that included 20 individual specimens of *Q. mongolica* or European *Q. robur*, respectively. Each of the *Q. robur/Q. mongolica* specimens were sampled from 10 geographically-widespread populations. To develop a panel of polymorphisms for North American white oaks, we sequenced chloroplast genomes from additional specimens representing the following species: *Q. alba, Q. bicolor* Willd., *Q. garryana* Douglas ex. Hook., *Q. lyrata* Walter, *Q. macrocarpa* Michx., *Q. michauxii* Nutt., *Q. prinoides* Willd., and *Q. stellata* Wangenh.

For marker validation, DNAs from 13 *Quercus* species were screened. *Q. mongolica* and *Q. dentata* Thunb. represented Asian oaks (Far East Russia) with 200 and 10 specimens, respectively. *Q. robur, Q. petraea* and *Q. pubescens* WillD. represented European oaks, with 360, 210, and 200 specimens, respectively. Finally, eight white oak species were screened from North America (*Q. alba, Q. bicolor, Q. garryana, Q. lyrata, Q. macrocarpa, Q. michauxii, Q. prinoides, Q. stellata*), with between 5 and 25 specimens per species.

### Next-generation sequencing analyses

Aliquots of DNA (∽0.5 - 1 μg) were sheared to a median length of ∽300 bp and converted into sequencing libraries using Illumina TruSeq v.2 kits at the USDA Forest Service (Corvallis, OR; individual samples, each indexed with dual-index adapters) or GATC Biotech AG (Konstanz, Germany; pooled samples). Sequencing was performed using three approaches: (A) using the Illumina MiSeq with 2x150 bp paired-end reads for individual *de novo* genome reference assemblies; (B) using the Illumina MiSeq with 2x300 bp paired-end reads for pooled samples and reference-guided mapping; and (C) using the Illumina HiSeq with 100 bp single-end reads for individual North American species and reference-guided mapping. All reactions used version 3 sequencing chemistry. Information on raw clusters, sequence yield, and approximate target sequence (chloroplast genome) coverage depth is provided in Table 1.

**Table 1:** Results of next-generation sequencing for oak reference assembly and polymorphism screening. Indexed individuals of oaks were sequenced using 150 bp paired-end reads and evaluated using *de novo* assembly (reference assembly). Pooled individuals were sequenced using 300 bp paired-end reads and evaluated using reference-guided assembly (polymorphism screen).

|  | Reference Assembly |  |  |  | Polymorphism Screen |  |
| --- | --- | --- | --- | --- | --- | --- |
|  | QUMO5 | QUPE2 | QURO2<sup>1</sup> | QUAL | QUMON Pool | QUROB Pool |
| Sequences | 6,183,339 | 5,281,517 | 6,676,925 | 4,280,049 | 17,370,324 | 28,438,226 |
| Bases sequenced (Gbp) | 1.747 | 1.502 | 0.977 | 1.207 | 10.42 | 17.06 |
| Proportion of reads mapping to cp genome | 3.3 | 1.9 | 7.6 | 0.4 | 0.95 | 2.25 |
| De novo contigs (length sum, Mbp) | 2,085 | 872 | 80 | 7,085 | -- | -- |
| Longest contig | 135,603 | 60,701 | 27,043 | 57,523 | -- | -- |
| Length chloroplast contigs | 4 | 12 | 19 | 20 | -- | -- |
| Variants relative to QUMO5 | 0 | 126 | 124 | 572 | 789 | 346 |

<sup>1</sup> *Q. robur* was sequenced using 100 bp single-end reads.

#### De novo reference construction

Raw read quality filtering was accomplished using Trimmomatic v0.30 (Bolger *et al.* 2014), and we removed reads with a mean Phred score less than 33. Reads were digitally normalized to a coverage of 20 using, khmer v0.7.1 (Crusoe *et al.* 2014), and kmer-filtered sequences were assembled using Velvet Optimizer v2.2.5 (https://github.com/Victorian-Bioinformatics-Consortium/VelvetOptimiser.git) and Velvet v1.2.10 (Zerbino & Birney 2008). K-mer lengths ranging from 21 to 121 were evaluated, with a final k-mer length of 121 selected for assembly. *De novo* contigs > 100 bp in length were screened for homology to the *Quercus rubra* chloroplast genome (NCBI NC_020152) using BLAT (Kent 2002). Contigs showing high similarity were retained for reference assembly and ordered against the *Q. rubra* chloroplast genome. Reference sequences were constructed to include the large single-copy (LSC) region, one of two inverted repeats (IR), and the small single-copy (SSC) region, equivalent to positions 1 −135,502 from *Q. rubra* NC_020152.

#### Reference-guided identification of SNP and indel variation

Reference guided read mapping and polymorphism detection was performed using CLC Genomics Workbench version 7.5.1 (CLC-bio, a Qiagen company; Aarhus, Denmark). The reference chloroplast sequence of the *Q. mongolica* individual QUMO5_CH_1 generated by *de novo* reference construction (see above) was used as reference for read mapping. The trimmed Illumina data of the two pools (Q. *mongolica, Q. robur*) and the trimmed HiSeq data of representative North American individuals were mapped to the reference scaffold using a length fraction of 0.9 and a similarity fraction of 0.94. Variants detected by CLC Genomics Workbench included SNPs and small indels, and these were exported to tab-delimited files and processed using an in-house script (*Variant Tools*, see below) to identify species-specific polymorphism.

### Post processing of identified SNPs and indels

To merge the SNP and indel tables and find common variants present in two or more individuals/pools, we developed *Variant Tools*, a command line program implemented in Ruby. This program merges individual sample SNP and indel tables (CSV format) produced by CLC GWB to create a multi-individual SNP and indel matrix. Required input options include the reference sequence (fasta format), an input directory containing the variant CSV tables, and an option specifying the input data type (SNP or indel). The reference fasta can contain one reference sequence or multiple reference contigs. Optionally, coverage tables (produced by CLC from read mappings to a reference) of every individual can be included in the analysis by specifying a directory containing coverage files (CSV format). Furthermore several filtering options are available to reduce the output according to user-provided thresholds. The output from *Variant Tools* is stored in a CSV file and contains several data columns: the reference sequence name, the reference position, the variant length, the calling type (SNP, MNP, deletion or insertion), the reference base(s), the alternative base(s) for every individual, the coverage for every individual at the reference position, different summary statistics, and sequences flanking the called variant.

The flanking sequences are calculated based on two given distance thresholds. An upper and a lower threshold define minimum distances in base pairs between two called variants on the genomic scale. If a variant occurs within the genomic range of the lower threshold no flanking sequence is created. If a variant resides within the genomic range of the upper threshold the length of the flanking sequence created is defined by the lower threshold. If no variant is found within the range of both thresholds a flanking sequence with the length of the upper threshold is calculated. The default thresholds are 75 bp and 50 bp.

The *Variant Tools* create different summary statistics while the variant matrix is generated. *The number of individual alleles deviating from the reference* is a count for all found variants in all individuals at a specific genomic position. *The number of alleles matching the reference with minimal coverage* is a count for all positions in all individuals where no variant has been called and that are supported by a minimum coverage. The threshold for the minimum coverage is specified by the user. The default threshold is set to a minimum coverage of 8. *Critical forward reverse balance* is an indicator for systematic sequencing errors and describes how many forward and reverse reads are supporting the called variant. The value is averaged over all individuals showing the variant.

The *Variant Tools* are open source software under ongoing development. They are available under the terms of the ICS license and can be obtained from https://github.com/ThuenenFG/varianttools.

### DNA extraction, PCR, restriction, and genotyping

*Leaves:* One cm^2^ of a single leaf was ground to powder in liquid nitrogen. Total DNA was extracted, following a modified ATMAB protocol by Dumolin *et al.* (1995). PCR reactions for leaf-derived DNA contained ∽30 ng template DNA, 10x PCR buffer, 1.5 or 1.75 mM MgCl_2_, 200 μM dNTPs, 0.4 unit AmpliTaq Gold DNA polymerase (ThermoFisher Scientific, Darmstadt, Germany), and 0.05 to 0.13 μM of each primer in a total volume of 15 μl. PCR was carried out in a Sensoquest Thermocycler (Göttingen, Germany) with a pre-denaturation step at 94°C for 10 min, followed by 25 to 30 cycles of 94°C for 45 sec (30 sec *trn*CD), suitable annealing temperature for each primer combination (between 52°C and 57°C) for 45 sec (30 sec *trn*CD), 72°C for 45 sec (1 min *trn*LF) and a final elongation at 72°C for 10 min. PCR amplification products were checked relative to a 100 bp ladder (Life Technologies, Martinsried, Germany) on a 1 % agarose gel stained with Roti-Safe GelStain (Carl Roth GmbH & Co. KG, Karlsruhe, Germany); afterwards, PCR products were run on an ABI3730 capillary sequencer. Fragment analysis was performed using GeneMarker^TM^ software v. 2.4.0 (Softgenetics, State College, PA, USA).

*Timber:* For genotyping analysis of timber-derived DNA, mostly a special DNA extraction protocol has been developed and patented (Lowe *et al.* 2015) based on the CTAB method. Exceptionally the innuPREP Plant DNA Kit from Analytik Jena (Germany) was used. Due to the small amount of total DNA extracted from timber, the DNA quantity wasn’t measured; rather, a standard dilution of 1:10 (DNA:water) was used for all PCR reactions. PCR conditions were similar to those used for leaf material, but with slight modifications (described in Results).

For one marker (*trn*DT), a restriction enzyme digest was used to reveal a SNP polymorphism in the amplified fragment. The restriction digestion reaction contained 10μl PCR product, 2ul 10x CutSmart^^®^^ buffer, 0.5μl enzyme (FastDigest^^®^^ *Hinf*I, New England Biolabs, Ipswich, MA) in a final volume of 20μl. The reaction lasted 15 min at 37°C followed by an inactivation at 80°C for 20 min. Restriction products were either visualized relative to a 50 bp ladder (Life Technologies, Germany, Martinsried) using an 8 % polyacrylamide gel stained with ethidium bromide, or using an ABI3730 capillary sequencer.

### Probabilities for fixation of gene markers

The Thünen-Institute of Forest Genetics possesses a large collection of reference samples that contains oak species from Europe (*Quercus robur, Q. petraea*), North America (*Q. alba, Q. macrocarpa*, and others) and Asia (*Q. mongolica, Q. dentata).* Based on the numbers in this collection of white oaks from different continents, we computed the maximal potential frequency of variants that were not observed using 95% confidence intervals (Newcombe 1998). This can be described as a method to determine the risk (potential error rate) that a genetic variant (allele/haplotype) assumed to be exclusive to one continent is found in one or more individuals originating from another continent. The calculations were carried out using the online forma at http://vassarstats.net/prop1.html based on 962 European, 325 Asian and 61 American white oak individuals for the gene markers psaI-ycf4, *psb*E-*pet*L, *trn*LF, and *trn*CD. For the gene marker *trn*DT the sample sizes were 115 European, 425 Asian and 19 American white oak individuals.

## Results

### Next-generation sequencing, reference genome assembly, and identification of cpDNA length variants in white oaks

Next-generation sequencing of four indexed oak exemplars (*Q. alba, Q. mongolica, Q. petraea. Q. robur*) yielded between 1.3 and 1.9 Gbp per individual (Table 1). *De novo* contig assembly with Velvet produced between 872 and 7,085 contigs, and the longest contigs from each assembly were from the chloroplast genome. Our best *de novo* chloroplast assembly derived from *Q. mongolica* QUMO5_CHI_1, which was represented by a single large contig totalling 134.6 kb, and it spanned the three main chloroplast genome regions (large single copy, inverted repeat, small single copy) in their entirety. Alignment of these contigs against the published *Q. rubra* chloroplast genome yielded an alignment of 136,475 nucleotides (alignment excludes one inverted repeat). Parsimony analysis of ordered *de novo* contigs and *Q. rubra* as an outgroup produced a topology similar to the topology previously established with chloroplast DNA restriction site analysis (Manos *et al.* 1999; Fig. 1), with the Old World white oaks *Q. mongolica, Q. petraea* and *Q. robur* resolving as a sister group to the New World *Q. alba.* Within this white oak chloroplast genome reference data set, we identified 672 single nucleotide variants and 189 indels (Fig. 1).

**Fig. 1:**
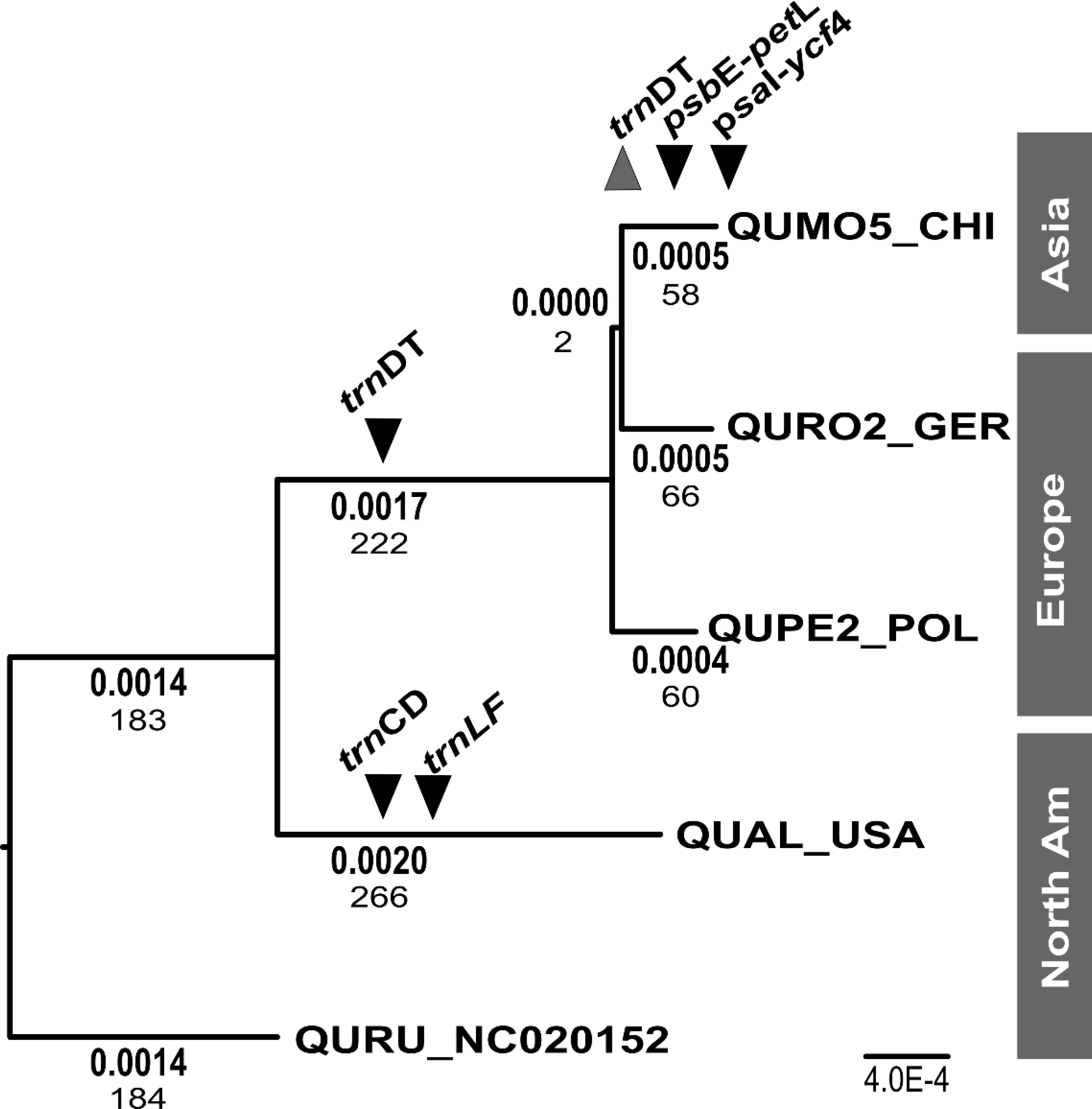
Phylogenetic relationship among chloroplast genomes of white oak species representing Old World and New World lineages. The best maximum likelihood tree is shown for four white oak chloroplast genomes (*Q. mongolica; Q. robur; Q. petraea; Q. alba*) and one outgroup genome (*Q. rubra).* Inferred branch lengths in maximum likelihood substitutions are shown in bold; the number of substitutions inferred from parsimony shows these values. The phylogenetic resolution of informative indel markers are shown in black inverted triangles, and the resolution of the diagnostic PCR-RFLP marker is shown as a grey triangle.

Next-generation sequencing of the *Q. mongolica* and *Q. robur* DNA pool produced 17.4 and 28.4 million paired-end 300 bp sequences (10.4 and 17.1 Gbp of sequence data respectively; Table 1). Mapping of paired-end reads from the *Q. mongolica* and *Q. robur* pools against the *Q. mongolica* QUMO5_CHI_1 chloroplast genome reference revealed 346 variant positions for the *Q. robur* pool, and 789 variant positions for the *Q. mongolica* pool (Table 1). After read mapping, the two variant tables were compared to filter those variants that showed fixed differences between the continents. Because the aim of this study was to develop markers that differentiate between the continents, the next step was to reduce the dataset to these variants appearing with a frequency between 95 and 100%. This analysis left five indels and 15 SNPs. For marker development, we focused in indels due to their simplicity of analysis. Checking of these indels within the mapping revealed a (T)N microsatellite, two indels with a difference in the fragment length of two bp, and one indel with one bp difference. These were removed from the further analyses and only the two longest indels in two spacer regions (*psaI-ycf4*, *psbE-petL*), one with four and one with six bp difference, remained.

Short read sequences from eight North American species were also mapped to the QUMO5 reference, and we specifically searched for indels differentiating North American species from the reference QUMO5 with a frequency of 95 to 100%. We identified three indels that consistently differentiated North America from Asia. These length polymorphisms were found in three spacer regions (*trn*LF, *trn*CD, and *trn*DT) and they ranged in length from 2 bp to 8 bp.

### Primer design, marker validation and resequencing

For the five indel-including cpDNA regions, primers were designed using the reference QUMO5 to amplify fragments ranging from 110 bp to 190 bp. A preliminary validation performed with three individuals each of *Q. robur, Q. petraea, Q. mongolica, Q. alba*, and *Q. macrocarpa* revealed that all seven cpDNA regions could successfully be amplified by PCR. Subsequent Sanger sequencing validated the sequence of the intervening region, the repeat type, and BLASTN analysis confirmed annotations (Table 2).

**Table 2:**
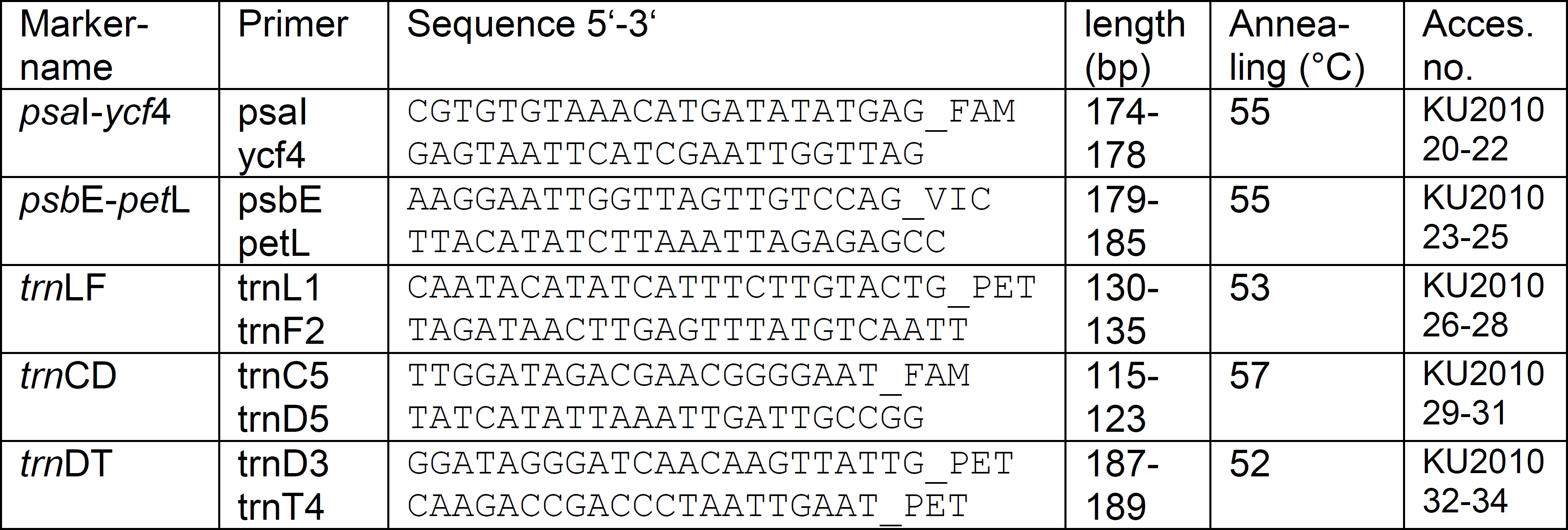
List of primers for the amplification and resequencing of the newly developed markers. Fluorescent-labeling of the primers is given in column “sequences”: FAM = blue, VIC = green, PET = red. In the last column, the accession numbers of the related markers for the three species *Q. robur, Q. mongolica* and *Q. alba* are given. “Length” means sequence length.

Two indels differentiate European (*Q. petraea, Q. robur*) and Asian (*Q. mongolica*) white oaks, and these are located in the *psaI-ycf4* linker (4 bp difference) and the *psbE-petL* linker (6 bp difference; Table 2), and these are inferred as mutations (deletions, specifically) are restricted to Asian white oaks. The further validation of these indels was conducted by screening the amplification products of 10 additional individuals of North American species, 50 individuals of two European species (*Q. robur, Q. petraea*), and two Asian species (*Q. mongolica, Q. dentata).* This validation revealed that white oaks from North America and Europe showed the same fragment length (Table 3), and confirmed that the Asian species shared deletions that resulted in shorter, diagnostic fragment lengths.

Three indels differentiate Old World (*Q. petraea, Q. robur, Q. mongolica*) and North American white oaks, and these are located in the *trn*L*-trn*F (*trn*LF) linker (5 bp difference), the *trn*C*-pet*N linker of the broader *trn*C*-trn*D (*trn*CD) linker region (8 bp) and in the *trn*E*-trn*T linker of the broader *trn*D*-trn*T (*trn*DT) linker region (2 bp; Table 2). These indels were validated with the same individuals of all above mentioned species. The validation revealed that white oaks from Asia and Europe showed identical fragment lengths (Table 3), and that the North American species shared mutations that resulted in diagnostic fragment lengths.

**Table 3:** Number of individuals per species and continent tested with the five markers, and fragment length based on sequencing for each marker and species.

| continent | species | <sup>1</sup> No. individuals | *psal-ycf4* | *psbE-petL* | *trnLF* | *trnCD* | <sup>2</sup> *trnDT* |
| --- | --- | --- | --- | --- | --- | --- | --- |
| Europe | *Q. robur* | 531 (103) | 178 | 185 | 135 | 123 | 86/101 |
|  | *Q. petraea* | 273 (12) | 178 | 185 | 135 | 123 | 86/101 |
|  | *Q. pubescens* | 158 (0) | 178 | 185 | 135 | 123 |  |
| USA | *Q. alba* | 15 (4) | 178 | 185 | 130 | 115 | 88/101 |
|  | *Q. macrocarpa* | 12 (7) | 178 | 185 | 130 | 115 | 88/101 |
|  | *Q. bicolor* | 7 (4) | 178 | 185 | 130 | 115 | 88/101 |
|  | *Q. garryana* | 4 (0) | 178 | 185 | 130 | 115 |  |
|  | *Q. lyrata* | 4 (2) | 178 | 185 | 130 | 115 | 88/101 |
|  | *Q. michauxii* | 5 (1) | 178 | 185 | 130 | 115 | 88/101 |
|  | *Q. stellata* | 8 (1) | 178 | 185 | 130 | 115 | 88/101 |
|  | *Q. prinoides* | 6 (0) | 178 | 185 | 130 | 115 |  |
| Asia | *Q. mongolica* | 316 (420) | 174 | 179 | 135 | 123 | 86/71 [30] |
|  | *Q. dentata* | 9 (5) | 174 | 179 | 135 | 123 | 86/71 [30] |

<sup>1</sup> The number of individuals tested includes numbers for all loci except *trnDT*, which is given in parentheses.
<sup>2</sup> Fragment lengths for *trnDT* show the lengths after restriction digestion with *HinfI*. Asian species contain an additional internal restriction site that yields one additional unlabeled fragment after *HinfI* digestion; these are shown in brackets.

Resequencing (Sanger) of the *trn*DT region identified a single nucleotide polymorphism (SNP) differentiating Asia from Europe and North America. The SNP lies within a *Hinf*I restriction site, and restriction digestion of this region was predicted to yield three fragments in Asian white oaks and two fragments in European and North American species (all oaks share one *Hinf*I site). By labeling the forward and reverse amplification primer, restriction digestion of the PCR fragment with *Hinf*I allows the visualization of two of the three fragments, one of which is diagnostic for Asian white oaks due to its truncated length. Since this single region offers the possibility to differentiate all three continents with one assay, we decided to include this SNP into the marker set. In total, the five markers have been evaluated with samples sizes ranging from 516 to 1078, with samples representing 13 oak species from the three continents (Table 3).

The nucleotide variation shown to be characteristic for Asian white oaks (*Q. mongolica* and *Q. dentata*) are also present in the complete chloroplast genome of one individual of *Q. aliena*, another Asian white oak species (GenBank accession KP301144.1; Lu *et al.* 2015). We computed the probabilities of fixation for the gene markers. Since we require a minimum of two independent markers for continent assignment, we performed the calculations for the risk of not identifying a rare variant in the reference samples based on a combination of two markers. By this means the risk of not identifying a rare variant in the reference samples was calculated to be less than 0.022 % for Europe, 0.0051 % for Asia and less than 0.098 % for America. Thus, it is extremely unlikely that studied gene markers are not fixed to just one variant in the different groups.

### Marker set design and optimization for timber

All above described analyses were performed using single PCR reactions and fragment analyses to optimize each marker separately. Subsequently, the markers were successfully multiplexed for fragment analysis using the fluorescence labeling as given in Table 2 (Fig. 2). For the development of the markers and multiplexing, DNA from fresh leaves was used. The protocol was later optimized for DNA from timber. From our experience, DNA from timber is more sensitive to all PCR parameters, thus, all markers were singly tested with DNA from timber and the PCR was optimized for the DNA from timber. Differences in the PCR conditions used for the two different materials are given in Table 4. Due to the sensitivity of timber DNA in PCR, multiplexing of the PCRs of the five markers is not advisable, but multiplexing of the markers on the sequencer worked as well with timber DNA as with DNA from leaves. The only difference is that the PCR product from leaf DNA is diluted 1:50 and the PCR product from timber DNA 1:10 for use on the sequencer.

**Fig. 2:**
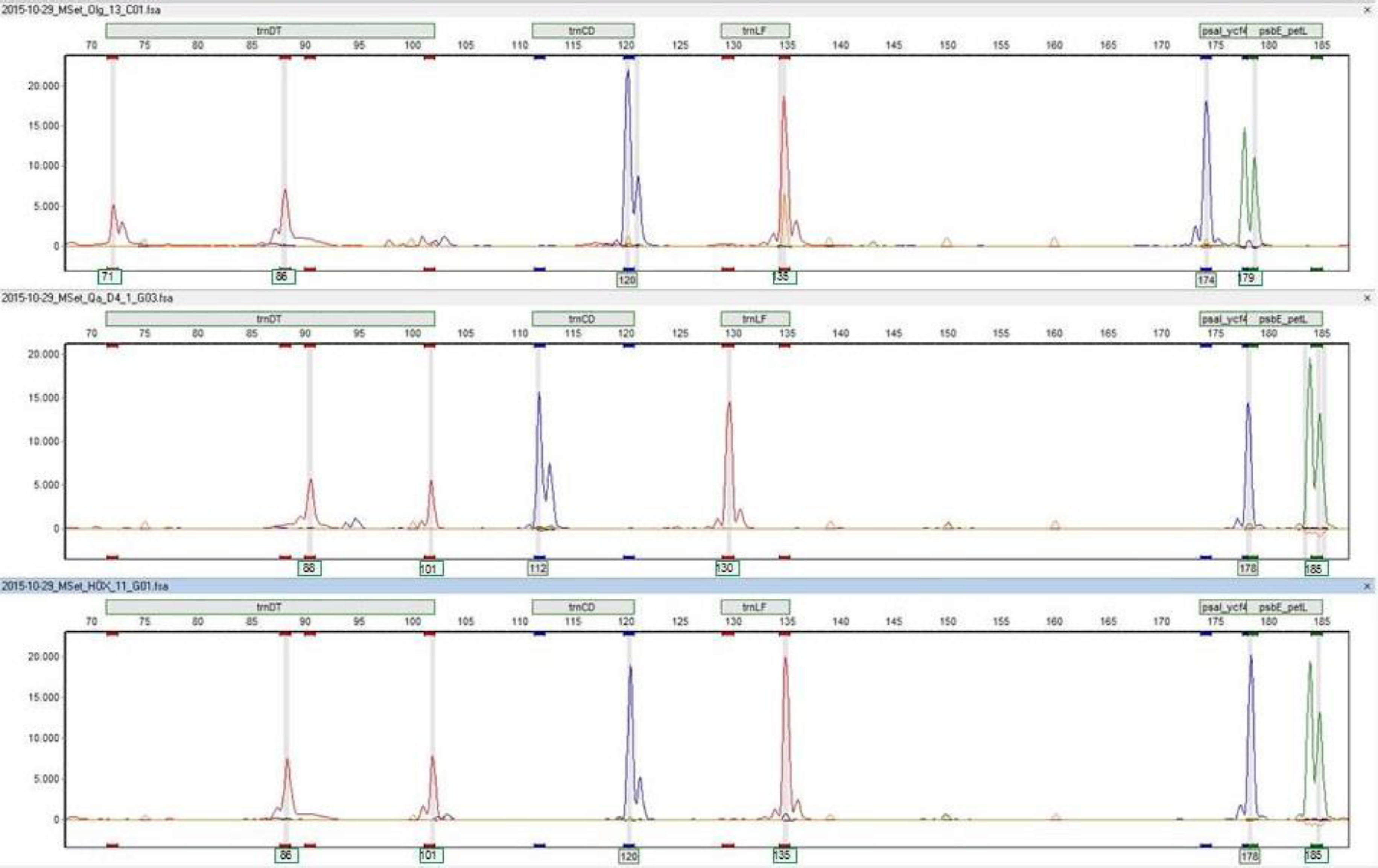
Fragment patterns of the five markers for individuals from Asia (top), North America (middle) and Europe (bottom). The sequence sizes for each peak as given in Table 3 are shown beneath the peaks. The first blue peaks appear smaller (112, 120) than the sequenced length (115, 123) given in Table 3. The color code of the peaks is as described in Table 2.

**Table 4:** PCR conditions compared for leaf and timber. Only the differences are shown, all other parameters are as given in material and methods.

|  | Leaf |  |  |  | Timber |  |  |  |
| --- | --- | --- | --- | --- | --- | --- | --- | --- |
|  | PCR cycles | Conc. MgCl<sub>2</sub> | Enhancer | Conc. Primer | PCR cycles | Conc. MgCl<sub>2</sub> | Enhancer | Conc. Primer |
| psal-ycf4 | 30 | 1.75 mM | no | 0.07 µM | 40 | 2.0 mM | no | 0.1 µM |
| psbE-petL | 30 | 1.75 mM | no | 0.05 µM | 40 | 2.0 mM | no | 0.05 µM |
| trnLF | 30 | 1.75 mM | yes | 0.2 µM | 40 | 2.0 mM | yes | 0.2 µM |
| trnCD | 25-30 | 1.5 mM | yes | 0.13 µM | 35 | 2.5 mM | yes | 0.3 µM |
| trnDT | 25 | 1.75 mM | yes | 0.1 µM | 40 | 2.0 mM | yes | 0.1 µM |

Our analyses used a capillary sequencer to visualize length polymorphisms of these fragments. However, due to the large size differences of these indels, all markers can be distinguished on a polyacrylamide gel, even for differences as small as two base pairs, as shown for the fragment *trn*DT (Fig. 3). In this way, polymorphisms can be screened in laboratories where no sequencer is available.

**Fig. 3:**
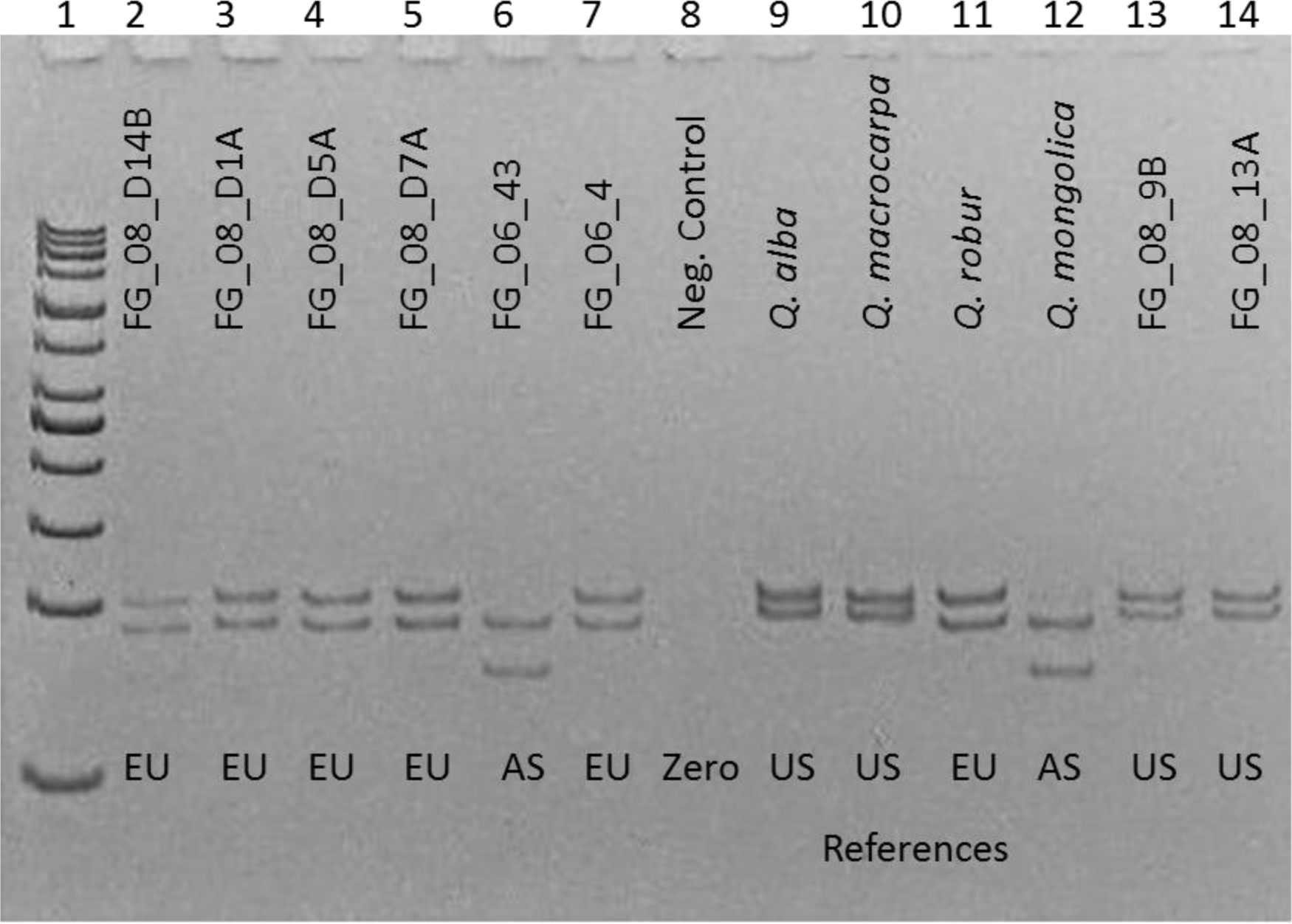
Marker *trn*DT visualized on a polyacrylamide gel. Lane 1: 50 bp ladder, lane 8: zero control, lane 2-7 and 13-14: analysis of wood-derived DNA, its location is inferred from genotypes, lane 9-12: references from North America (US), Europe (EU) or Asia (AS), respectively.

The functionality of the optimized PCR protocols and the multiplexed sequencer runs has been tested by means of orders worked on in the “Thünen Centre of Competence on the Origin of Timber”. Timber from these orders included highly processed wood as flooring or parquet, as well as treated solid wood samples as different parts of furniture, barrels or boards, and unused solid wood as firewood. In total, over 80 processed timber samples and 130 treated solid wood samples have been evaluated (data not shown). Based on our experiences so far we have a success rate of sufficient DNA amplification for our gene markers for 58 % for solid wood samples.

## Discussion

A set of five chloroplast markers have been developed and optimized to analyze DNA from timber to identify the continental origin of white oak wood. Small fragment sizes (< 200 bp) were chosen because genotyping success with DNA from timber is highest when fragments under 200 bp are targeted. This has recently also been shown for DNA from old and dried insect specimens of museum collections when using mitochondrial barcoding regions (Mitchell 2015). For the identification of haplotypes within oak species from wood samples, Deguilloux *et al.* (2003) similarly developed chloroplast microsatellite and SNP markers that targeted small DNA fragments.

The sequencing data revealed no specific indels to differentiate oak species within the classical barcoding regions *mat*K, *rbc*L or the linker *trn*H-*psb*A (e.g., Chase *et al.* 2005; 2007; Lahaye *et al.* 2008; Hollingsworth 2011; Tripathi *et al.* 2013). Recently, the barcode regions matK and *trn*H-*psb*A were evaluated for their power to discriminate select species from oak sections *Cerris, Heterobalanus* (= “Group Ilex”), *Lobatae*, and *Quercus* (Simeone *et al.* 2013). In this study, the matK region proved to have too low resolution for the differentiation within the genus; interestingly, for *trn*H*-psb*A, the variability was too high to identify fixed interspecific differences (Simeone *et al.* 2013).

The intergenic linkers *trn*LF, *trn*CD and *trn*DT we found valuable in this work have been widely tested in population and evolutionary genetic studies of plants, and they show wide variation in their ability to discriminate species and lineages (Shaw *et al.* 2005). For example, the intergenic linker *trn*LF proved to be not variable enough for overall barcoding approaches (Chase *et al.* 2005). For the trial of phylogenetic reconstructions this *trn*LF linker lacked variation in closely related species (Dong *et al.* 2012). Similarly, differentiation within the genus *Populus* failed using *trn*LF (Schroeder *et al.* 2012). Nevertheless, there are other examples for successful use of this marker in molecular systematics (Martin-Bravo *et al.* 2007; Pirie *et al.* 2007 and citations therein) and for unraveling of the phylogeny of different plant species (Drábková *et al.* 2004). Similarly, *trn*CD and *trn*DT have been used in comprehensive studies of chloroplast DNA diversity in European white oaks (Bordács *et al.* 2002; Cottrell *et al.* 2002; Csaikl *et al.* 2002; Fineschi *et al.* 2002; Jensen *et al.* 2002; König *et al.* 2002; Olalde *et al.* 2002; Petit *et al.* 2002a, b, c). In Japan, four oak species have been differentiated using *trn*DT among other chloroplast markers (Kanno *et al.* 2004). Hence, as for many regions within the chloroplast, the applicability of these spacers to questions of species identity depends on the specific phylogenetic and geographic context where they are used.

Forensic applications of molecular markers are already established with regard to illegal wildlife trade (e.g. parrots: Goncalves *et al.* 2015; sea turtles: Rehman *et al.* 2015) or for identification of products made of endangered animal species (e.g. ‘whale meat’: Baker *et al.* 2010; horn: Yan *et al.* 2013). The barcode of wildlife project (http://www.barcodeofwildlife.org/) has been originated especially for this purpose. Further on control of the seafood market in different countries is well supported by usage of barcoding markers (e.g. Pappalardo & Ferrito 2015; Carvalho *et al.* 2015). For identification of illegal logging of tropic tree species the use of molecular markers is already widespread (e.g. Höltken *et al.* 2012, Nithaniyal *et al.* 2014, Hartvig *et al.* 2015), but the methods are less established for tree species from temperate zones. Thus, the presented markers should be applied to give commercial vendors of white oak wood the possibility to exercise ‘due diligence’ when placing timber on the European market and the public authorities to control timber imports should questions emerge on the correct declaration of wood.

## Acknowledgements

We are grateful for the financial support by German Federal Ministry of Food and Agriculture (BMEL), the Deutsche Bundesstiftung Umwelt (DBU) and the USDA Forest Service International Programs Office. We thank Lasse Schindler for developing of the timber specific DNA extraction protocol, and Aki M. Höltken for initial discussions. We are grateful to Vivian Kuhlenkamp, Laura Schulz, Susanne Jelkmann and Ann-Christine Bergmann for technical assistance. We also thank collaborators and institutions that provided samples of North American oaks, including the Morton Arboretum, Lisle, IL (Andrew Hipp, Marlene Hahn), North Carolina State University, Raleigh, NC (Paul Manos), and the USDA Forest Service (Paul Berrang, Andy Bower; Britton Flash; Steven Forry; Ben Gombash; Mark Twery). Finally, we thank Mark Dasenko, Matthew Peterson, Chris Sullivan (Oregon State University Center for Genome Research and Biocomputing) for assistance with sequencing, and Brad Kinder, Alex Moad, and Shelley Gardner (US Forest Service) for project and planning assistance.

## Data Accessibility

DNA sequences from Sanger sequencing of the indel containing fragments derived from this study have been deposited in GenBank (Accession numbers: KU201020-KU201034). Next-generation sequences supporting this study are available from the NCBI GenBank as BioProject PRJNA269970 (SRA Accessions SRS954648, SRS954649, and SRS954650). SRA accession numbers of the next-generation sequencing data of the two DNA-pools are coming soon. Draft chloroplast genome references will be submitted to Dryad soon. Genotype calls for samples will be available on the EVOLTREE website (http://www.evoltree.eu/index.php/e-recources/databases) soon. The open source software *Variant Tools* is available on https://github.com/ThuenenFG/varianttools.

## Author Contributions

Y.Y. coordinated sample collections in Russia and China, Y.Y. and H.S. coordinated sample collections in Europe, and R.C. coordinated sample collections in North America. H.S. and B.K. performed the analysis of sequence pools, identified variants, and designed indel screens. H.S. coordinated the Sanger sequencing and the genotyping. R.C. and T.J. constructed libraries for North American species, constructed de novo genome references, and assisted with variant analysis. B.D. did statistical calculations and together with R.C. initiated the project. M.M. developed the Variant Tools. H.S., B.K., B.D. and R.C. were involved in overall coordination. H.S. wrote the manuscript and all authors contributed to and approved the final manuscript.

